# Decoding heart failure subtypes with neural networks via differential explanation analysis

**DOI:** 10.1101/2025.03.03.641151

**Authors:** Mariano Ruz Jurado, David Rodriguez Morales, Elijah Genetzakis, Fatemeh Behjati Ardakani, Lukas Zanders, Ariane Fischer, Florian Büttner, Marcel H. Schulz, Stefanie Dimmeler, David John

## Abstract

Single-cell transcriptomics offers critical insights into the molecular mechanisms of heart failure with reduced or preserved ejection fraction. However, understanding these mechanisms is hindered by the growing complexity of single-cell data and the difficulty in unmasking meaningful differential genes signatures among heart failure types. Machine learning, particularly deep neural networks, address these challenges by learning transcriptional patterns, reconstructing expression profiles and effectively classifying cells but often lacks interpretability. Recent advances in explainable AI (XAI) offer tools to clarify model decisions. Yet pinpointing differentially regulated genes with these tools remains challenging.

In this study, we introduce a novel method to identify differentially explained genes (DXGs) based on importance scores derived from custom-built neural networks. We highlight the superiority of DXGs in identifying heart failure subtypes-specific pathways that provide new insights into different types of heart failure. Offering a robust foundation for future research and therapeutic exploration in expanding transcriptome atlases.

## Main

Despite the implementation of secondary preventative therapies cardiovascular diseases remain the leading cause of morbidity and mortality in the aging society^1^. Reliable biomarkers are crucial for early detection of heart failure and facilitate timely diagnostics and personalized treatment strategies. However, the complexity of the underlying transcriptomic alterations, compounded by the different pathophysiological mechanisms and inter-individual heterogeneity of each disease, is one of the major challenges while detecting biomarkers or understand and distinguish certain disorders like different types of heart failure (HF)^2,3^. Large-scale single-cell and single-nucleus transcriptomic sequencing (scRNA-seq and snRNA-seq, respectively) of clinical patients or relevant mouse models are widely used to discover biomarkers and understand disease mechanisms that contribute to hypertrophic HF induced by high blood pressure aortic stenosis (AS), HF with reduced ejection fraction and with preserved ejection fraction (HFrEF and HFpEF, respectively).^4–9^

Recent advancements in computational power, and machine learning approaches have offered an unbiased framework to analyse scRNA-seq data^10^. Despite the noise in large transcriptome data caused by data sparsity, biological variability, technical artifacts, and sequencing errors^11,12^, computational methods have shown accurate performance on a variety of tasks, such as, cell type annotation^13^, imputation^14^ and batch aware data integration^15^. A particularly successful machine learning approach is the denoising autoencoder (DAE), that learns a low-dimensional latent space through non-linear transformations and defines a decoder to reconstruct the initial expression of genes per cell^16^. In addition, deep feed-forward neural networks demonstrated capability of precisely classifying transcriptomic data into various biological categories and disease treatments^17,18^.

Biomarker or novel therapeutic targets are often identified from scRNA-seq data by differential gene expression (DGE) analysis methods. This typically employs a variety of statistical approaches to address challenges such as sparsity, dropout events, and overdispersion. The analysis includes nonparametric methods such as the distribution-independent rank-sum test^19^ or the classical parametric t-test^20^. Another approach is the use of hurdle models, which separately account for zero-inflation and gene expression levels, enhancing sensitivity in sparse data^21^. Additionally, likelihood ratio tests (LRT), implemented in various methods^22^, use generalized linear models (GLMs) based on Poisson or negative binomial distributions, or logistic regression models predicting group membership based on each feature individually. Other notable approaches include pseudo-bulk methods, where single-cell data is aggregated within biological replicates to approximate bulk RNA-seq data. This enables the use of bulk RNA sequencing tools for a robust DGE analysis, while accounting for biological variability^23^.

These analyses often use nonparametric statistical tests or tests based on negative binomial distributions to identify significant differences in expression between healthy and diseased conditions^24,25^. However, their statistical frameworks often lack the capacity to capture more complex patterns of disease- and cell type-specific gene expression changes^26,27^.

Taking advantage of DAEs and feed-forward neural networks we devised an approach for the identification of heart failure related specific targets and biomarkers by detecting disease-relevant patterns. Our neural network was designed to accurately classify patient and mice snRNA-seq data from healthy and diseased hearts enabling it to assign each cell a multiclass label that describes its species, cell type, and disease state. In addition, we suggest a new statistical approach to investigate which gene expression patterns were important for the model’s decision using Shapley values^28^, an approach from explainable AI (XAI). Shapley contribution scores have recently been used in combination with scRNA-seq data to study regulation at the gene-level^29^ and for validation purposes for computational methods regarding cell type annotation in murine cardiac hearts^30^. Here we advance this methodology, by designing a statistical test tailored for Shapley gene contribution scores from neural networks, which provides an alternative approach for biomarker identification using a concept we term differentially explained genes (DXGs). We provide evidence that DXG derived biomarkers are more reliable than results from classical DEG methods.

## Results

### Neural network designed to classify cells by species, cell type, and heart failure state

We designed a neural network to process normalized snRNA-seq datasets from diseased and healthy ventricular samples from both patients and mouse models, which were unified into a large tabular format. To ensure that only representative nuclei were used in our training, validation, and test sets; we rigorously filtered, retaining only high-quality nuclei (Methods, Extended Data Fig. 1). We excluded nuclei with an inadequate number of reads or high mitochondrial content, thereby reducing inaccuracies and biases that could potentially affect the model’s ability to learn class-specific signatures. After filtering the biological samples, we obtained the following nuclei counts: 144,677 from healthy patient hearts, 40,205 from healthy mouse hearts, 41,689 from patients with hypertrophic hearts caused by AS, 3,594 from mice with transverse aortic constriction (TAC), 19,330 from patients with HFrEF, 7,046 from mice after 28 days following permanent ligation of the left anterior descending artery (LAD) to induce HFrEF, 1,712 from patients with HFpEF and 13,827 from the double hit murine HFpEF model^31^.

In order to integrate gene expression counts from human and mouse, we restricted the analysis to one-to-one orthologs between the human and mouse genomes computed using OrthoIntegrate^32^. This dataset included annotations for species, cell type, and disease state, which were encoded in the neural network to be learnt as individual classification problems (Fig. 1a). We ensured that our integration and automated cell type annotation was correctly capturing cell clusters by investigating the UMAP per species (Fig. 1b-c, Extended Data Fig. 2c). By inspecting the expression of published cell type specific markers^5,33^ (Fig. 1d-e), we re-annotated the automated annotation (Extended Data Fig. 3) into cardiomyocytes (CMs), endothelial cells (ECs), fibroblasts (FBs), immune cells (ICs), neuronal cells (NCs), pericytes (PCs) and smooth muscle cells (SMCs).

**Fig.1:**
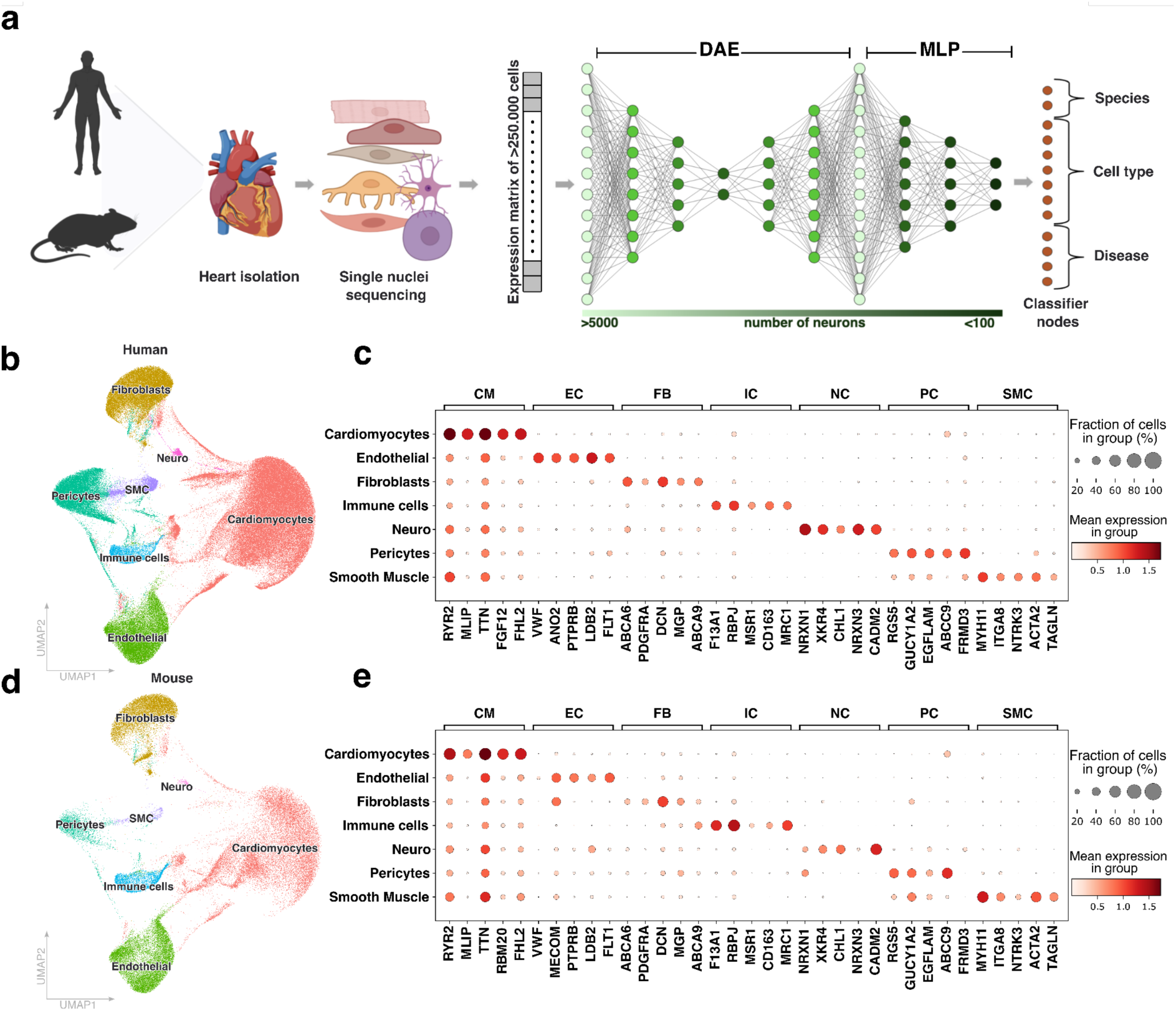
Cell type annotation based on marker genes for deep learning models. **a**, Schematic overview of workflow. Single nuclei RNA-seq data is obtained from hearts of either diseased or healthy humans and mice. The expression matrix undergoes initial preprocessing before being passed to a denoising autoencoder (DAE) for learning a streamlined representation focused on a narrow bottleneck of the transcriptomic data. Afterwards this representation is used to train a multilayer perceptron, enabling the classification of cells based on species, cell type, and disease state. **b,d**, Uniform manifold approximation and projection of cell types identified in **b)** human and **d)** mice. **c,e**, Average expression and fraction of cells expressing known marker genes for cardiomyocytes (CM), endothelial cells (EC), fibroblasts (FB), immune cells (IC), neuronal cells (NC), pericytes (PC) and smooth muscle cells (SMC) in **c)** human and **e)** mice.

The neuronal network is composed of two components. First, the DAE processes the normalized reads and learns based on a symmetrical layer structure. By reducing the mean squared error loss (L2-loss), a biologically and technically denoised representation of our HF data was generated^16^. Second, a multi-layer perceptron (MLP) classifier uses the autoencoder’s denoised gene expression values for each cell as input to classify cells into three distinct classes, differentiating the 2 species, 7 cell types and 4 disease states. To compensate for imbalanced labels, we defined a custom F1-Loss function based on the harmonic mean of precision and recall. For testing purposes, we reserved one complete biological sample from each species and disease state, guaranteeing a robust performance assessment on hold-out data.

### Optimization of neural network architectures and performance comparison for predicting species, cell type and heart failure types

In order to have a baseline performance we used a standard logistic regression (LR) classifier for the three classes and evaluated its performance (Extended Data Fig. 2a). Classification rates on hold-out data showed that the LR classifier model poorly captured the high dimensional transcriptomic data (F1-score: 0.6615) showing the need for a more complex classification approach (Extended Data Fig. 2b,d,f).

We conducted extensive parameter tuning of the neural network, enhancing performance and allowing subsequent biological interpretation of the autoencoder and the MLP components by exploring the layer’s neuron counts using a Hyperband bandit-based algorithm (Fig. 2a-b; Extended Data Fig. 4; Methods).

**Fig.2:**
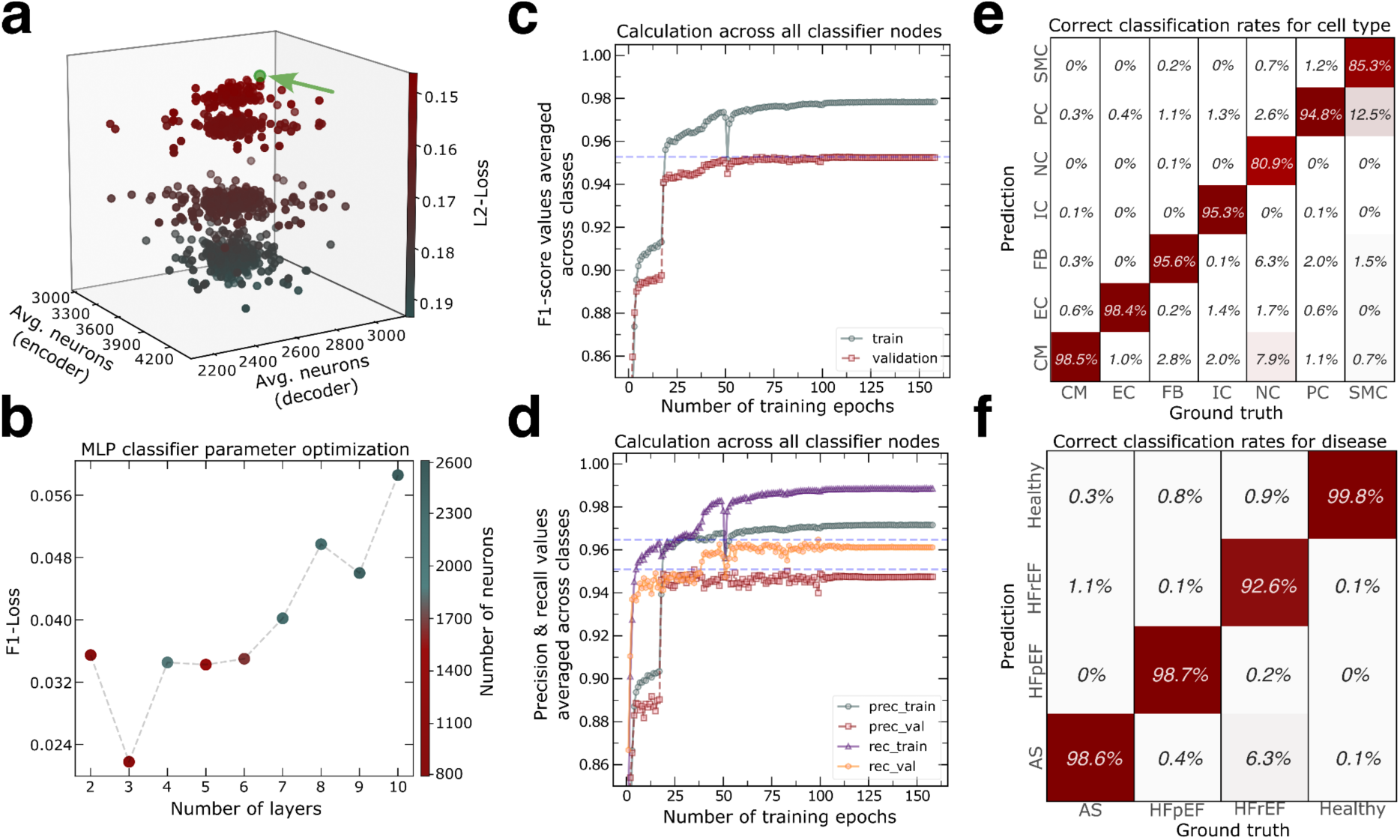
Hypertuned neural network reaches high accuracies predicting heart failure. **a**, Three-dimensional scatter plot illustrating the average number of neurons in the decoder and encoder layer of the DEA, along with the corresponding L2-Loss for each set of parameters tested during hyperparameter tuning. **b,** Highlighting the lowest F1-Loss values during the hyperparameter tuning of the MLP classifier for each layer configuration. Color indicates the total number of neurons in the network. **c**, F1-Score calculations per epoch of the MLP classifier’s performance averaged across all classes. **d**, Precision and recall calculations per epoch of the MLP classifier’s performance averaged across all classes. **e-f,** Column wise calculations on percentages of correctly classified samples for the classifiers output nodes regarding **e)** cell types and **f)** disease state.

After evaluating the results, we decided to use 5000, 2400, and 350 neurons for the three layers in the encoder, including the latent space, and 350, 2200, 5150 neurons in the decoder. While symmetric autoencoders are common practice, studies indicate that an asymmetric neuron design could yield improved performance^34,35^. Therefore, we performed all analyses using the best-performing autoencoder after hypertuning (L2-Loss: 0.1393, Fig. 2a; Extended Data Fig. 4b). Further hyperparameter tuning for the MLP classifier architecture revealed that a three-layer structure achieved the lowest F1-Loss value (F1-Loss: 0.0217, Fig. 2b; Extended Data Fig. 4c). Consequently, we used this three-layer network in combination with the best-performing DAE for all subsequent analyses.

By averaging across all classes, we obtained an overall F1-score of 0.9784 (Fig. 2c) for training data and 0.9528 for validation data, indicating a class-unbiased and accurate model structure suitable for downstream model interpretation. To address the potential issue of certain classes being underrepresented during model training, we calculated F1-scores separately for species, annotated cell types, and HF conditions. (Extended Data Fig. 4d-f). In the validation set, we observed notably high scores across all three groups: approximately 1.00 for species identification (Extended Data Fig. 4d), 0.9562 for cell type classification (Extended Data Fig. 4e), and 0.9789 for disease diagnosis (Extended Data Fig. 4f). Additionally, we determined if there was a significant difference in precision and sensitivity among the three groups that could impact the F1-score calculation. Therefore, we conducted a separate analysis to examine the precision and recall values (Extended Data Fig. 4g-j). We observed high precision for both metrics, suggesting that the model achieves strong performance by accurately predicting classes with few false positives, while maintaining high sensitivity and thus minimizing false negatives.

Furthermore, we evaluated the individual misclassifications between categories in the prediction of hold-out samples. To achieve this, we calculated confusion matrices that accurately captured the correct classification rates for each of the 13 attributes. On average our model achieved 100%, 92,7%, and 97,4% accuracy for species, celltype and disease classification, respectively (Fig. 2e,f; Extended Data Fig. 2e). These high classification rates among all classes highlight the models capability to accurately distinguish between mouse and human cells for each cell type and disease state allowing an in-depth biological interpretation of the MLP classifier decisions.

### XAI analysis using Shapley values identifies genes that serve as predictors for species, cell types and heart failure conditions

In order to shed light on the importance of individual genes that contribute to classification performance we estimated local attribution scores for each single cell measurement and each class using Shapley values (Fig. 3a). This approach used the gene expression of 200 cells randomly taken from each cell type in each biological sample. We obtained a set of Shapley values corresponding to the explained data for each of the classification categories covering 52,925 cells and 16,545 genes per category.

**Fig.3:**
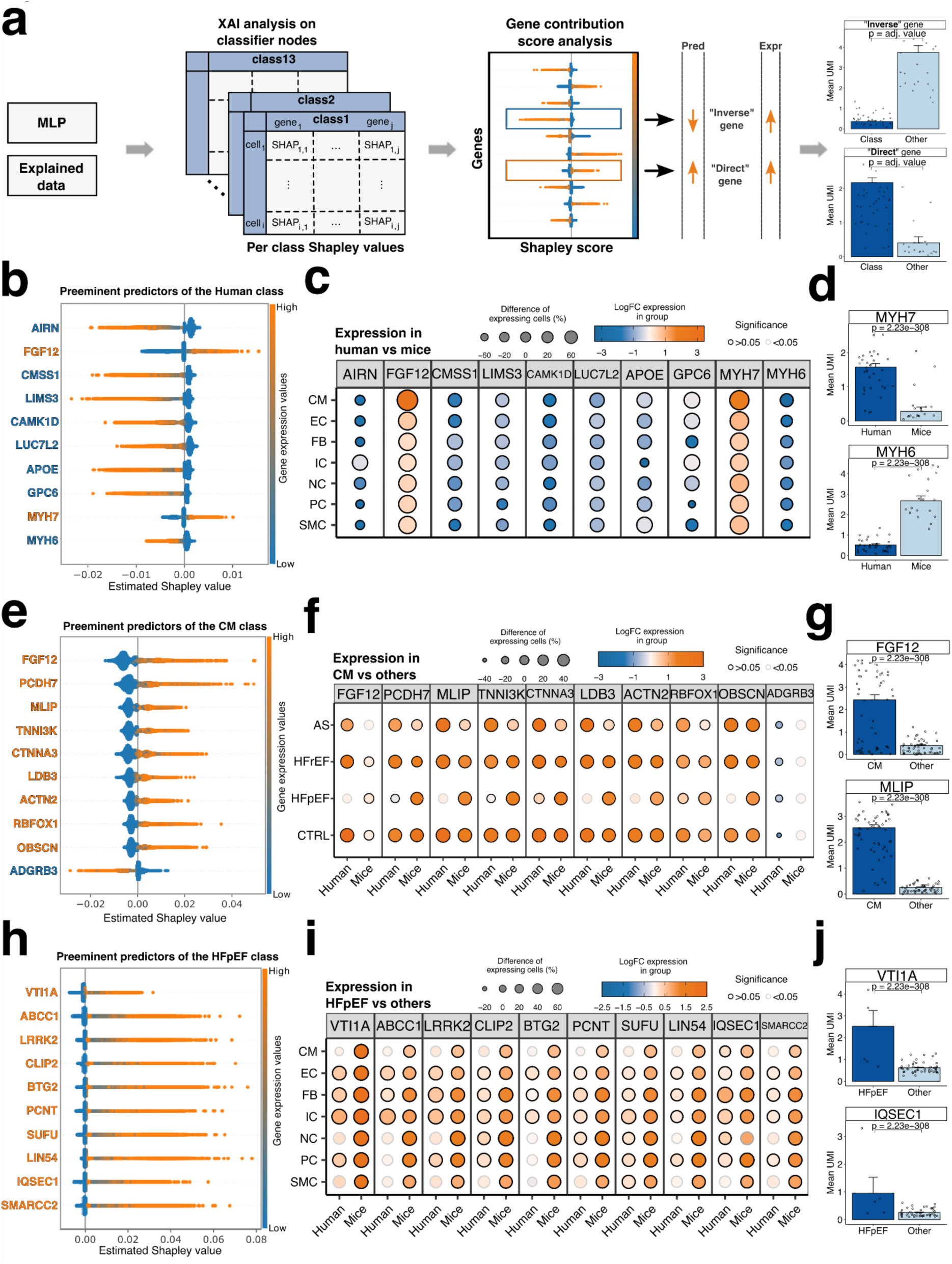
XAI analysis reveals known and potential markers relevant for species, cell types and heart failure. **a**, Schematic overview of Shapley values obtained by analyzing the hidden layers of the MLP classifier. Each classifier node produces a set of values corresponding to the gene’s contribution in that class. Prediction and expression are directly correlating or following an inverse relationship based on the type of predictor. Inverse markers show negative Shapley values and are associated with an absence of expression, while direct marker expressions are correlating with positive values. **b,e,h,** Ten most contributing predictors visualized by a beeswarm plot for **b)** human cells, **e)** CMs and **h)** HFpEF cells. An orange gene name indicates a positive correlation between expression and prediction for that class, while a blue gene name indicates a negative correlation. **c,f,i,** Dot plot illustrating the log fold change in gene expression and the percentage change of cells expressing the indicated genes. Panels represent the following **c)** human predictors by cell type across all disease states, **f)** CM predictors by disease and species and **i)** HFpEF predictors by cell type and species. **d,g,j,** Averaged unique molecular identifiers (UMI) for two selected genes for **d)** human cells against mice cells, **g)** CMs against all other remaining cell types, **j)** HFpEF cells against the remaining conditions. Statistical significance was determined using the Wilcoxon test on single cell expressions.

The Shapley values for the human and mouse categories exhibit contrasting patterns owing to the binary nature of the decision task, where only one of two outcomes can be realized. Consequently, gene predictors contributing to one of them will obtain opposing scores for the other category. For example, high Shapley values for the myosin heavy chain genes *MYH7/Myh7* were identified as key predictors for differentiating between human and mouse cells (Fig. 3b), showing a direct correlation with its high expression levels in human cells (Fig. 3c-d). Conversely, genes *MYH6/Myh6* exhibited an inverse relationship with the classifier’s predictions, where high expression levels of *MYH6/Myh6* corresponded to lower prediction values for cells associated with humans. Notably, when examining the classification node representing mouse cells, we observed opposite outcomes (Extended Data Fig. 5a, Fig. 3d) consistent with the known difference in myosin heavy chain switch in humans versus mice^36,37^.

Additionally, we found that the long non-coding RNA, *AIRN*, exhibited a pattern similar to that of *MYH6*. High expression levels were associated with cells from mice hearts, while low or absent expression corresponded to cells from the human patients (Fig. 3b-d). Indeed, *AIRN* has been identified as a regulator of the insulin-like growth factor receptor IGF2R gene, which constitutes the largest known imprinted domain in mice, but not in humans^38,39^. Successful capture of these ratios demonstrates that the model accounts for well-established differences between human and mouse cell expression patterns, which suggests its capacity to explore relevant gene markers between both species.

Furthermore, we investigated the relevant predictors for the model’s decision on cell type annotation. In our analysis, we identified several highly relevant predictor genes in CMs, including the fibroblast growth factor *FGF12*, the cardio-muscular interacting MLIP, and the TNNI3 interacting kinase *TNNI3K* (Fig. 3e). Each of these genes are well-known markers for CM identification. Additionally, we discovered other top predictor genes that serve as valuable markers for CMs identification. For instance, high expression levels of the protocadherin family member 7 (*PCDH7*) and the cell-cell adhesion protein *CTNNA3* were crucial for identifying CMs (Fig. 3e). On the other hand, the model was able to capture down-regulated markers. For instance, the absence of the secretin receptor family member *ADGRB3* was a significant predictor (Fig. 3e). We verified that the predictors’ Shapley values match the gene expression found in the underlying transcriptomic data (Fig. 3f-g). Analysis of the predictors relevant for identifying the remaining cell types identified well-established marker genes commonly used for cell annotation (Extended Data Fig. 5b-g). Similar to the findings in CMs, we identified among the top predictors some previously uncharacterized genes that could potentially aid in the detailed annotation of the identified cell type populations. Notably, *ST6GALNAC3* emerged as a key predictor with high relevance in ECs, while the absence of *LAMA2* also played a significant role in their characterisation. In contrast, in fibroblasts, the high expression of *MEG3* and *COL8A1* was crucial for classification (Extended Data Fig. 5b-c).

### Using Shapley values reveals novel candidate HFpEF biomarkers

Cardiac-specific biomarkers are crucial for distinguishing between the different HF states, allowing the stratification of patients into risk categories for early diagnosis and targeted intervention. We applied Shapley value analysis on the classifier nodes to predict whether a cell corresponds to HF resulting from AS, HFrEF or HFpEF (Extended Data Fig. 6, Fig. 3h-j). We could identify several novel biological candidates across cell types that were crucial for predicting HFpEF. For instance, the gene *VTI1A* emerged as a key predictor for cells in HFpEF, representing a novel candidate for patients with this condition. This gene encodes for a protein from the N-ethylmaleimide-sensitive factor attachment protein receptor (SNARE) family and has been implicated in cell signalling via endo-lysosomal trafficking in neurons^40,41^. While its role in the heart is understudied, one study has linked *VTI1A* variants to association with length of the QRS complex, an ECG-based measurement of the beating heart, in UK Biobank participants. QRS prolongation was previously associated with poorer prognosis in HFpEF^42,43^. *ABCC1*, also referred to as *MRP1*, is a membrane-bound transporter that facilitates the extracellular efflux of both endogenous and exogenous molecules from cells, playing a key role in cellular detoxification and drug resistance mechanism^44–46^. Notably, *ABCC1* mediates the removal of cyclic GMP-AMP (cGAMP), an important activator of the innate immune response^45^. In mice, elevated cGAMP levels were observed to impair vascular remodelling, and endothelial cell proliferation via dephosphorylation of YAP1^47^. Furthermore, heightened cGAMP serum concentrations have been observed in HF patients^48^, reinforcing the potential importance of this candidate.

*LRRK2* encodes a multi-domain protein involved in intracellular processes like microtubule dynamics and vesicular trafficking^49^. While mutations in *LRRK2* are well-known in Parkinson’s disease^50^, its role in HFpEF is largely unexplored. However, a recent study reported a negative correlation between circulating levels of *LRRK2* with NT-proBNP, a biomarker for heart failure^51^. In mice, *LRRK2* expression was found to be upregulated in cardiac endothelial cells under HFpEF conditions^52^. Furthermore, *LRRK2* knockout mice subjected to TAC exhibited improved cardiac function and reduced adverse remodelling compared to their WT counterparts. These beneficial effects were found to be mediated through autophagy-related processes^53^. Although other studies have also confirmed a role for *LRRK2* in monocyte adhesion to endothelial cells via NF-κΒ^54^, a distinct requirement for the activation of sterile inflammation in HFpEF^55^. Each of these novel molecular targets could lead to more effective detection of HFpEF, with implications for improving disease prognosis and management.

### Shapley values can be used to conduct differential explanation analysis for identifying dysregulated genes

We then considered a method to statistically test whether Shapley values between two groups of cells show significant variations. Since these values were derived from transcriptomic data that reflect the gene expression values, we employed them to identify dysregulated genes among groups of interest, similar to traditional DGE analysis. To achieve this, we applied a Z-transformation to the Shapley values for each cell. This normalization step ensured that the relevant changes were comparable between different cells. Using the original labels per cell, we performed Student’s t-test to identify significant differences in Shapley values per gene in each group (Fig. 4a, Supplementary Table 3-6). Since we were no longer calculating DEGs based on expression levels alone, but instead using XAI, which reflects the predicted importance derived from the neural network, we refer to these genes as differentially explained genes (DXGs). To evaluate whether our lists of DXGs capture different sets of genes compared to the traditional Wilcoxon test on expression data, we first assessed the overlap and differences between DXGs and DEGs identified in CMs from HFpEF patients (Fig. 4b, Supplementary Table 7). Using a t-test on Shapley values comparing healthy-labeled CMs to HFpEF-labeled CMs we identified 2,198 dysregulated genes. In contrast, the Wilcoxon test on expression values identified 3,212 regulated genes. Of these, 1,178 genes were common to both methods (Supplementary Table 3&7). These findings demonstrate that each approach identifies unique genes that may be relevant biomarkers for HFpEF, although there is a significant overlap between the DXGs and DEGs.

**Fig.4:**
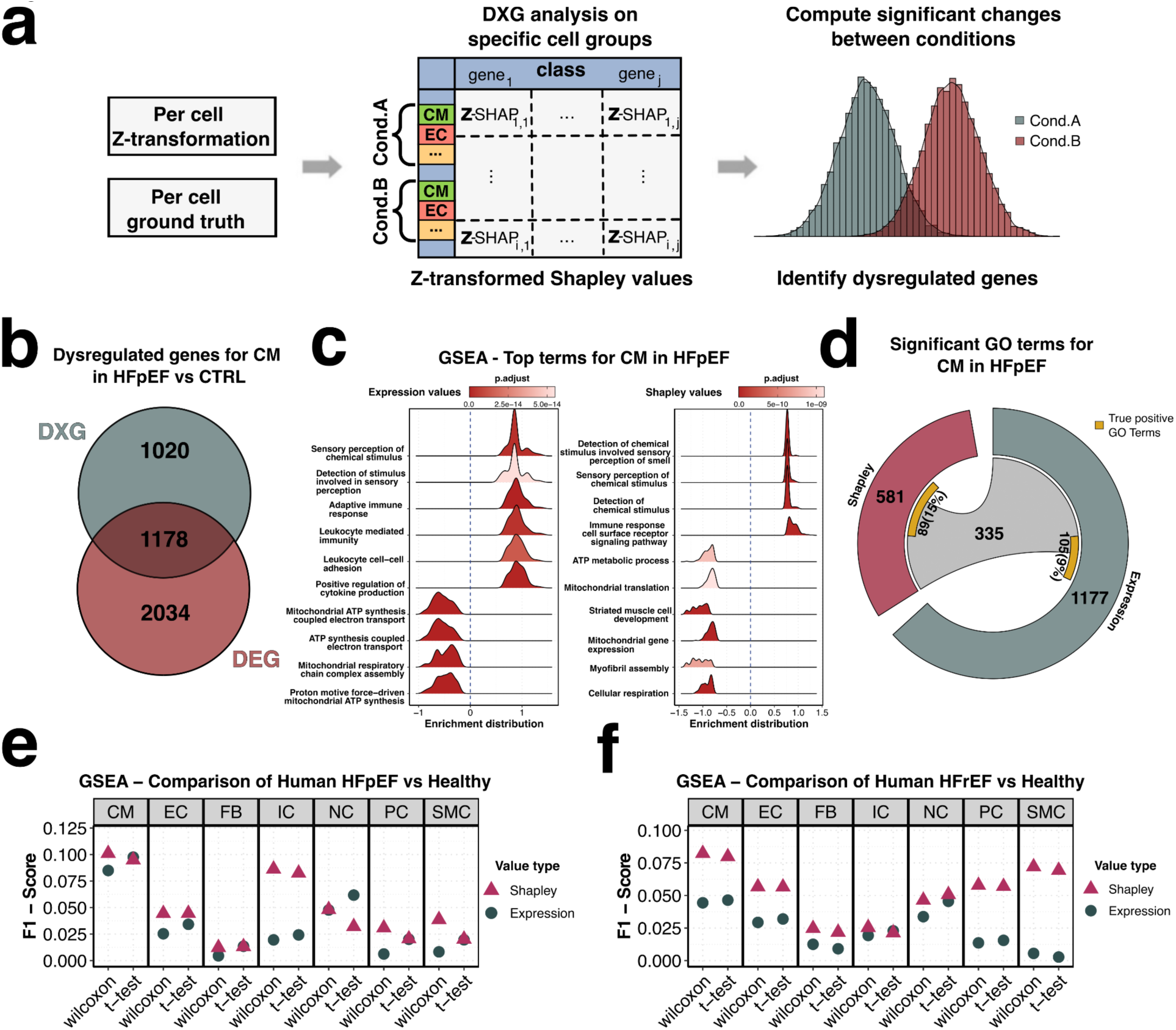
Exchanging expression values for Shapley contribution scores enhances the results of pathway analysis. **a,** Overview of the process for DXG analysis using Shapley values. First, the contribution scores are Z-transformed for each gene and combined with the ground truth data to identify relevant groups of cell types and conditions for comparison. The Shapley value sets for both conditions are then statistically tested to determine differences and identify dysregulated genes between the conditions. **b,** Comparison of the overlap of dysregulated genes identified in CM under the HFpEF condition versus the CTRL condition. DXGs were calculated by applying a t-test to Z-transformed Shapley values and DEGs were calculated through a Wilcoxon test applied on expression values. **c,** Top ten GSEA terms for CM under the HFpEF condition ranked by enrichment score, based on statistics from Wilcoxon test using expression values (left) and using the t-test applied to Shapley values (right). P-values were corrected using the Benjamini-Hochberg correction. **d,** Significant GO terms for CM under the HFpEF condition identified through GSEA analysis are shown for Shapley values (red) and expression values (grey). A chord highlights the terms shared between both analyses. Additionally, a yellow box displays the number of true positive GO terms for HFpEF in CM, illustrating the proportions found uniquely and commonly by each method. **e-f,** F1-Score calculations for GSEA results, performed separately using the Wilcoxon test and the t-test, are shown for each cell type. These calculations are based on expression values and Shapley values in **e)** the HFpEF condition and **f)** the HFrEF condition.

Next, we investigated whether the differences in identified genes led to the detection of distinct mechanisms associated with HFpEF in CMs. To explore this, we performed gene set enrichment analysis (GSEA) using p-values calculated for each gene with both methods (Fig. 4c, Methods). While both approaches highlighted similar significant terms, such as those related to sensory stimuli and ATP metabolic processes, DXG analysis also revealed additional terms relevant to CMs under HF conditions. Notably, one of the most regulated terms identified through this method pertained to muscle cell development (Fig. 4c). We then assessed the differences in identified pathways by comparing the significant pathways obtained from both methods and referencing them against a predefined set of true positive pathways known to be dysregulated in HF-affected CMs (Fig. 4d, Supplementary Table 8). The GSEA based on Shapley values identified 581 significant pathways, while the analysis using expression values yielded 1,177 terms, with 335 pathways overlapping between the two methods (Supplementary Table 9). Comparing these pathways to our true positive list, we found that 15% of the terms identified with Shapley values matched the predefined pathways, whereas only 9% of the terms from expression values matched. This indicates that while both approaches detect comparable numbers of true positive pathways for HF-affected CMs, the expression-based results are more prone to noise and the inclusion of irrelevant terms, potentially due to the sparsity of expression data (Fig. 4d-f).

To determine whether Shapley values consistently outperform expression values, we conducted GSEAs using both value types in combination with Wilcoxon and t-tests, repeating the analysis across all cell types and under both HFpEF and HFrEF conditions (Fig. 4e-f, Supplementary Table 9-10). We then calculated F1-scores by comparing the identified pathways to our previously defined list of true positives. The results demonstrated that Shapley values consistently outperformed expression values, further emphasizing their superiority in detecting dysregulated genes and their associated pathways.

## Methods

### Study samples

Detailed ethical information is published in the previously submitted manuscripts^4,5,32^. All other human tissue protocols were approved by the University Hospital of Mainz. Patients provided informed consent in compliance with the Declarations of Helsinki.

Human endomyocardial biopsies were collected from the left ventricle (LV) of patients at the University Hospital Mainz. HFpEF was defined as the presence of heart failure symptoms, echocardiographical or histological evidence of structural heart disease and an ejection fraction ≥50% in accordance with the current guidelines of the European Society for Cardiology and the American Heart Association^56,57^. Biopsies were processed for snRNA-Seq as previously described^4^. Briefly, samples were snap-frozen immediately after collection and stored at 80°C until further processing. Nuclei were isolated as previously described^4^ using a mechanical glass homogenizer in hypoosmolalic lysis buffer, separated from debris using an BD FACSAria Fusion Sorter and subsequently washed. After visual inspection of integrity, nuclei were subjected to snRNA-Seq library preparation using the 10X Genomics Chromium GEM3 v3.1 platform and sequenced on an Illumina NovaSeq 6000 (GenomeScan, Leiden, The Netherlands). Data on patients’ baseline characteristics can be found in Supplementary Table 2. Additionally, published human single-nuclei RNA-seq datasets were utilised in this study. Data for the healthy human myocardium were sourced from the PRJEB39602^5^ (Human Cell Atlas). The heart tissue was obtained from deceased transplant organ donors who were between 45 and 70 years old and showed unremarkable cardiovascular history. Data from patients with aortic stenosis were retrieved from E-MTAB-11268^4^. A detailed cohort description can be found in the respective publication. The HFrEF samples were obtained from the previously published E-MTAB-7869 dataset^32^.

This study adhered to the regulations for animal experimentation, and all procedures were approved by the relevant authorities. We obtained left ventricular tissue from TAC mice with procedures performed as previously described^32,58^, with an experimental endpoint at 28-days. Nucleus isolation and single-nuclei RNA-sequencing library preparation followed the guidelines outlined in Nicin et al.; NCVR 2022^4^. Healthy mouse heart dataset was sourced from our previous studies, available on the Array Express Data Portal under E-MTAB-7869 and E-MTAB-13264. These also contained the datasets for the HFrEF model. We also integrated mice treated with HFD/L-NAME from Kattih et al.^52^ available under the accession number E-MTAB-14589. Baseline information for mice model data can be accessed in Supplementary Table 2.

Single-nuclei transcriptomic sequencing results were preprocessed using CellRanger software (10x Genomics; v7.0.0). The initial phase involved demultiplexing and processing raw base count files using the implemented *mkfastq* tool. Reads from humans and mice were mapped to reference genome hg38 (GRCh38-2020) and mm10 (GRCm38-2020), respectively, utilizing CellRanger count with the option for including introns. Secondary data analysis was conducted using the programming language R (v4.3.1) and Seurat (v4.3.0)^59^. Barcodes with low (< 300) or high number of genes (> 6000) were filtered out and not considered for model training or data analysis purposes. In addition, barcodes exceeding a mitochondrial content of over 5% were discarded. Normalization is done by dividing the gene counts for each cell by the total counts for that cell and then multiplied by a scaling factor of 10,000. The result is then natural-log transformed and further clustering steps as well as the canonical correlation analysis integration of datasets were conducted following the guidelines provided from Seurat. We calculated one-to-one orthologs between human and mouse samples using the algorithm provided in the R package OrthoIntegrate^32^ and replaced mouse gene symbols with the human nomenclature. For further analysis, we were provided with 16,545 uniquely found genes between both species. Genes with multiple mappings were excluded, ensuring that only uniquely mapped orthologs were included for model training (Supplementary Table 1). The gene expression analysis and model training were conducted on this modified Seurat object structure. For annotation purposes, CellTypist (v1.6.3)^13^ was used, with the Healthy_Adult_Heart (v1) model specified for semi-supervised labeling of cell types using the provided reference for cells in healthy adult human hearts. Additionally, we carefully re-annotated misclassified clusters with the algorithm manually based on known markers, ensuring a high quality cell annotation. Cell clusters were visualized using UMAP dimension reduction implemented in Seurat and the average expression of cell type marker genes can be appreciated in the dotplot function implemented in the Python (v3.10.8) package Scanpy (v1.10.1)^60^.

### Creating training, validation and test data for model training

For every heart condition in patients and mouse models, we held back one patient and mouse for evaluation purposes. These samples were not used for any training on machine learning algorithms, ensuring no data leaks between training, validation and test sets. Training and validation sets were created using the normalized expression matrices of the remaining samples. Here, we randomly selected 80% of these remaining cells for defining the training set and the other 20% of cells were used for validation purposes while training. We ensured that cell types are equally presented in terms of percentages in all three datasets by using the R package caret (v6.0-94) and their implemented function for creating stratified datasets. The resulting sets were exported into Python for model training, validation and evaluation as well as for approaches of XAI.

### Dimensionality reduction and denoising autoencoder

The cell classification algorithm comprises two machine learning models developed using TensorFlow’s (v2.13.0)^61^ framework in the Python programming language. The first part describes the architecture of an autoencoder for dimensionality reduction and denoising purposes. Here, we sought to mitigate the inherent noise of the snRNA data and learn a low-dimensional representation by creating a symmetrical layer structure containing an encoding and decoding framework, with a narrow bottleneck layer. We performed hyperparameter optimization using Keras Turner’s Hyperband^62^ bandit-based algorithm and obtained a three-layer structure for the encoding part with 5000, 2400, 350 neurons for the first, second and latent space layers, respectively. Furthermore, we were provided with a two-layer structure for our decoding component with 5150 and 2200 neurons. We explored the first layer’s neuron count within a 2,000-neuron range centered around the initial value of 5,000, yielding a range of 4,000 to 6,000 neurons. For the second layer, we set a range of 1,000 neurons around 2,500, resulting in a range of 2,000 to 3,000 neurons. In the latent space, we defined a range of 200 neurons around 400, resulting in a range of 300 to 500 neurons. For the step size between neurons, we specified 50 neurons per layer. In addition, we train the hyperband algorithm to a maximum of 100 epochs per trial with a batch size of 10,240. For stochastic-gradient based optimization, we used the popular Adam optimizer while training. Neuron activation was conducted using the rectified linear unit activation (ReLU) function. A common choice for an autoencoder’s loss function is the mean squared error loss, also referred to as the L2-loss:

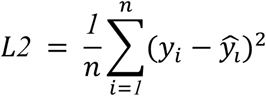

Here, *n* corresponds to the number of cells, *y_i_* corresponds to the predicted gene expression for cell *i* and *y*^_*i*_ to the actual gene expression for cell *i*. While training, we minimized this L2-loss and monitored the performance. If there were no improvement after 25 training epochs, the learning rate of the Adam optimizer will be reduced by a factor of 0.1 until a minimum learning rate is reached of 1×10^−7^. The optimizer was initialized with a learning rate of 0.001. The autoencoder was trained until 500 epochs or stopped after 50 epochs if there was no further improvement.

### Multilayer perceptron for multiclass classification

The expression matrices were reconstructed using the previously described autoencoder architecture and passed to the second part of the cell classification algorithm, which comprises the structure of a MLP multiclass classifier. This classifier returns for each cell the probability of belonging to a given species, cell type and disease group. As previously described, we performed hyperparameter tuning to define the optimal architecture and were provided with three layers containing 795, 230 and 105 neurons. The classification layer of the MLP consists of 13 neurons (classes), which determine probabilities individually using the sigmoid activation function on calculations of previous layers. Neuron activations located in hidden layers were conducted with the ReLU activation function. Labeling of cells was in a one-hot encoded manner, where 0 stands for a negative result for the given class and 1 for a positive one. The probability values for the 13 classes were assigned a value of 1 or 0, depending on whether they were above a threshold value of 0.5. We defined our macro F1-loss function as follows:

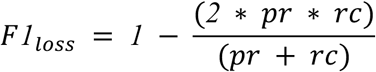

Here *pr* stands for the precision calculated by:

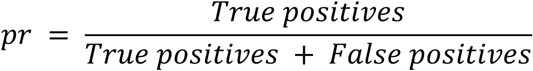

And *rc* represents the recall which was calculated by:

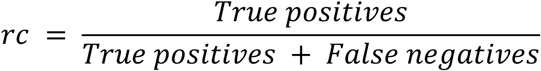

where *True positives* represents the number of correctly classified samples in that class, *False positives* the number of wrongly associated samples in that class and *False negatives* the number of samples that were not classified into a particular class despite belonging to it.

By minimizing this macro F1-loss function we were guaranteed to reach a maximum F1-score. The F1-score is a well established metric for scenarios where imbalanced classes are given and since cell types *in vivo* data follow such an imbalance, common accuracy calculations resulted in poor performance for underrepresented cell types. Considering the harmonic mean of precision and recall provides a balanced measure that penalizes extreme values and is therefore more suitable for our *in vivo* data. We carefully monitored all named metrics for training and validation datasets. As previously stated, we used the Adam optimizer with the option to reduce the learning rate after 25 epochs of no improvement and we introduced a maximum of 500 epochs or an early stop option after 50 epochs of no improvement.

### Evaluation of model performance

After finalizing the training procedure for the cell classification algorithm, we carried on with evaluating the performance of the classifier. For this, we predicted for each cell in our retained samples species, cell type and disease and compared them with the previously collected actual labels. We calculated a correct classification rate for each of the 13 classes independently. Additionally, we performed a confusion matrix calculation for the species, cell type and disease categories and visualized the results using the R package cvms (v1.6.2). This allowed us to inspect if the model performs adequately on unseen data for all major and minor groups of class combinations.

### Gene contribution score calculations using Shapley values

For calculating gene contribution scores for all 13 classes, we made use of Shapley values using the SHAP Python package^63^ (v0.44.0), which allows the approximate calculation of these on deep learning models. Based on the training dataset, 1,000 cells were randomly selected as the background distribution for calculations. DeepExplainer was used as the preferred explainer given the dimensionality of the data and the shape of the model. The explainer was initiated with the MLP classifier and the previously defined randomly selected background and used to calculate gene contribution scores for cells in the test dataset. We excluded mitochondrial genes from being possible predictors. Additionally, we standardized the scores of genes per cell by applying a rowwise Z-score transformation as follows:

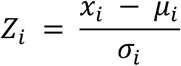

Here *x_i_* corresponds for the Shapley value for a given gene of cell *i*, μ*_i_* is the average of all Shapley values for all genes in cell *i*, and *σ_i_* is the standard deviation of the Shap values for cell *i*. We took advantage of the beneficial data structure of anndata^64^ objects and converted each of the thirteen sets of z-transformed Shapley values into an individual object. We then proceeded with concatenating them into one data structure keeping track of their individual class assignment.

### Identifying differentially regulated genes based on Shapley values

Given the data structure of AnnData objects and the ability to add metadata, we proceeded to add ground truth information for each cell based on species, cell type and disease state. With this additional knowledge, we can trace each cell directly back to its original label and thus define groups of interest that we want to compare. Considering the rather novel nature of Shapley values, we conducted a Shapiro-Wilk test on gene values per cell to test if they follow a normal distribution. We used the Scipy Python package (v1.11.1) for accessing the shapiro function and conducting the cell-by-cell test for normality. The resulting p-values were adjusted for multiple testing using the Bonferroni correction method. We proceeded with defining a function, which selects a set of cells with Shapley values corresponding to a class of interest. Every original cell in the test dataset is therefore selected with z-standardized Shapley values for a specific class. Given the ground truth data stored in the metadata of the AnnData object and the actual ground truth data file, we defined groups of cells corresponding to a given species, cell type and disease state and compared their gene values with a second group of cells of interest for significant changes in their z-standardized Shapley values. We introduced two possible ways of testing by implementing a Wilcoxon rank-sum test for values that do not follow a normal distribution and a Student’s T-test for samples that follow normality. The p-values were adjusted by false discovery rate (FDR) using the Benjamini-Hochberg correction. The fold changes between classes were transformed by taking the average cube root of the value per gene following this equation:

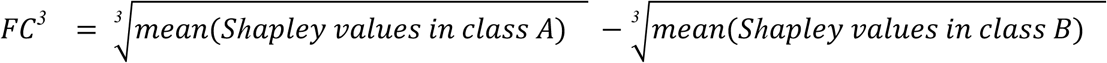

This calculation is conducted for each gene found in both class *A* and class *B*, similar as in traditional DGE analysis. The cube root transformation was defined for negative, zero and positive values, which can be present in the Z-standardization of data. In addition, it reduces the right skew similar to a logarithmic transformation to base 2.

### Differential gene expression analysis and GSEA analysis

We performed DGE analysis using the R package Seurat. For species, cell type and disease state specific results, we used the previously created Seurat object and downsampled by only containing cells which will be compared. We used the FindMarkers function with default parameters for filtering gene lists for log2foldchange and Bonferroni corrected p-values. We used Wilcoxon and MAST tests implemented in Seurat. For gene set enrichment analysis (GSEA) we performed DGE analysis with non filtered gene lists. The score per gene for the analysis was calculated by:

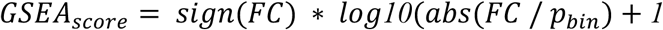

with *FC* representing the fold change of the given gene and *p*_*bin*_ representing a binned p-value, which is set to 0.1 if it was lower than 0.05 or otherwise set to 1. This encourages a weighted balance between p-values and fold changes, without disregarding the change in expression of the given gene. The GSEA was performed by using the fast gene set enrichment (fgsea) method implemented in clusterProfiler R package (v.4.13.2).

### Benchmarking DGE analysis results

We identified a set of gene ontology (GO) terms for each cell type that are known to be regulated in heart failure^4,5^ (Supplementary Table 8). These terms were then compared to the results of the GSEA analysis based on the DGE analysis using Seurat’s Wilcoxon and t-tests, as well as the list of DXGs using Wilcoxon and t-tests. Finally, we calculated an F1-score using the found terms from our predefined set as true positives and the remaining terms which are not in the defined sets will be treated as true negatives. All terms which were not found were used as false negatives. We illustrated the GSEA performances using expression and Shapley values in a dotplot created with the ggplot2 package.

## Discussion

In this study, we present a novel approach combining a DAE with a MLP and an adjusted loss function to classify species, cell types, and specific heart diseases based on single-nuclei transcriptomic data. This strategy demonstrates the sufficiency of individual nuclei transcriptomes for detecting various heart failure types (HFpEF and HFrEF) in mice and humans. Our model achieved F1-scores of > 0.95 on validation data for the classification of cell types and disease states.

When combined with XAI tools, specifically Shapley values, these models have the potential to significantly enhance unbiased biomarker discovery in cardiovascular research. In this pilot study, our model captured complex transcriptomic patterns, such as species-specific expression of myosin heavy chain genes (*MYH7* and *MYH6*), thus validating the model’s biological interpretability. Importantly, this analysis reaffirmed additional known interspecies differences while uncovering novel insights, such as the role of *AIRN* in distinguishing mouse and human samples, underscoring the potential of our approach for identifying key markers of cardiac phenotypes.

Shapley values have first been introduced by Lundenberg et.al^63^ and since then have been used to predict cell type markers from single-cell data in early developmental stages^65^, identify severe risk factors for cardiovascular diseases^66^ or pinpoint disease associated enhancers^29^. Here, our multiclass MLP model uniquely demonstrates that Shapley values can identify biomarkers for cell types, species, and diseases from a single MLP model that was trained with scRNA-SEQ data.

Our exploration of hidden layers enabled the identification of predictive genes for species, cell types, and heart failure (HF) conditions. Novel findings, such as the implication of *VTI1A* and *ABCC1* in HFpEF, provide valuable leads for further research, especially given their links to processes like vesicular trafficking and immune regulation, which may play a crucial role to drive immunological alterations in heart failure^67,68^. Furthermore, our model effectively captured established cardiac markers, while revealing potential novel markers, such as *CTNNA3* and *PCDH7* for cardiomyocytes, thus advancing cell type annotation and phenotypic stratification.

By calculating DXGs, we proposed a novel method for identifying dysregulated genes and biomarkers. In contrast to differentially expressed genes, which only consider the gene expression under two conditions, the comparison of Shapley values considers all possible permutations of genes and ranks their importance across all possible subsets of genes. We hypothesize that this approach reduced the number of irrelevant DEGs potentially caused by data sparsity or other confounding factors. Yielding more targeted and reliable disease-specific markers. Shapley values or similar contribution scores in general could provide a robust alternative to traditional statistical methods by integrating the advantages of machine learning techniques to overcome the limitations of traditional DEG analysis, particularly in sparse and noisy datasets.

Despite the promising results, several limitations should be acknowledged. Processing large transcriptomic datasets required substantial computational power, which may be challenging without access to GPU servers. Additionally, the small dataset, particularly for human samples, constrained the scope of our findings and may affect the generalizability of the results. Additionally, restricting the dataset to orthologous genes between humans and mice potentially excluded relevant heart failure-related genes, which may limit the depth of biological insights^32^.

Efforts to balance the classifier achieved reasonable fidelity but highlight the need for further refinement to reduce potential biases in model predictions. The reliance on randomly selected background datasets for Shapley value calculations may introduce inaccuracies, necessitating future efforts to refine these calculations with more representative data. Although our neural network outperformed logistic regression, newer machine learning models, such as transformer-based approaches pre-trained on extensive collections of datasets with a masked training objective, are already showing promising results in single-cell transcriptomics and may achieve superior performance, warranting further exploration in future studies^69^.

Gene set enrichment analysis (GSEA) on real-world data may not fully capture the intricacies of heart failure conditions, requiring further experimental validation. Additionally, imbalances and sparsity in the dataset pose risks of misclassification, underscoring the need for model improvement and expanded datasets.

To address these limitations, future efforts should focus on increasing sample sizes, incorporating more diverse datasets, and exploring emerging machine learning frameworks. Expanding the model to include additional genes and leveraging more representative background datasets for Shapley value calculations could further enhance its accuracy and applicability. Especially the future integration and training with already established large organ cell atlases^5,70,71^ could further prove the robustness and the clinical importance of the genes detected by our established method. Yet, integrating experimental validation will be crucial to confirm the biological relevance of identified markers and improving translational potential.

The introduction of DXGs as a concept adds a powerful dimension to traditional differential gene expression analysis. By demonstrating a higher signal-to-noise ratio and increased precision in identifying pathways relevant to HF phenotypes, our approach highlights the added value of incorporating machine learning-based feature attribution in transcriptomic studies. This methodology may outperform traditional expression-based analyses in uncovering biologically relevant pathways. Future studies are essential to address if the identified genes using this novel approach indeed have biologically validated functions to ultimately document the value of the developed tool.

In conclusion, our work underscores the potential of advanced neural network architectures integrated with explainable AI to enhance our understanding of cardiac biology. The ability to reliably classify cell types and disease states while identifying key predictive genes and pathways positions this framework as a transformative tool in cardiac research. Future studies could leverage this approach to investigate other complex biological systems, ensuring broader applicability and deeper insights into disease mechanisms.

## Data availability

The patient healthy heart data used in this study can be obtained through the Human Cell Atlas (HCA) data explorer under the project name “Cells of the adult human heart” (https://explore.data.humancellatlas.org/projects/). For our analysis, we included only tissues from the left ventricle and septum to match the region of our obtained disease samples. The patients with the donor id: H0015, H0020, H0025, H0026, H0035, H0037, CBTM-362C, CBTM-364B, CBTM-390C, CBTM-417C, CBTM-423C and CBTM-473C were used as controls. We excluded samples from donor CBTM-361B and CBTM-386C since they were diagnosed with hypertension and diabetes, respectively. We obtained patient data diagnosed with HFrEF from the ArrayExpress database using the accession number E-MTAB-13264. Additionally, we collected under the latter mice HFrEF samples and three healthy mice (n1-n3). The remaining healthy mice were collected from the ArrayExpress database using the accession number E-MTAB-7869. The snRNA-seq dataset from hypertrophic hearts caused by aortic stenosis, used in this study, was obtained from Nicin et al.^4^ and is available in the ArrayExpress database under the accession identifier E-MTAB-11268. We also collected three mice treated with HFD/L-NAME from Kattih et al.^52^ available under the accession number E-MTAB-14589. The datasets generated and analyzed during this study are available in the ArrayExpress repository under accession number E-MTAB-14753. This includes data from two human HFpEF patients and three TAC mice. We provided a summary containing baseline characteristics under Supplementary Table 2.

## Code availability

The Python and R code used to generate the models and the analysis can be accessed through the github repository at https://github.com/MarianoRuzJurado/RuzJurado_et_al_2024.

## Acknowledgements

The study was funded by DZHK (German Centre for Cardiovascular Research), partner site RheinMain, Frankfurt am Main, Germany to D.J., German Research Foundation (DFG), SFB 1531 - Project Nummer 456687919, project B01 to SD, project S03 to MHS; German Research Foundation (DFG), SFB 1366, Project Number 394046768; and the Dr. Rolf M. Schwiete Stiftung, Projekt 08/2018 to S.D. We are grateful to the Engelhardt lab at the Institute of Pharmacology and Toxicology at Technical University of Munich (TUM) for contributing left ventricular samples from mice subjected to TAC.

---

**Extended Data Fig.1:**
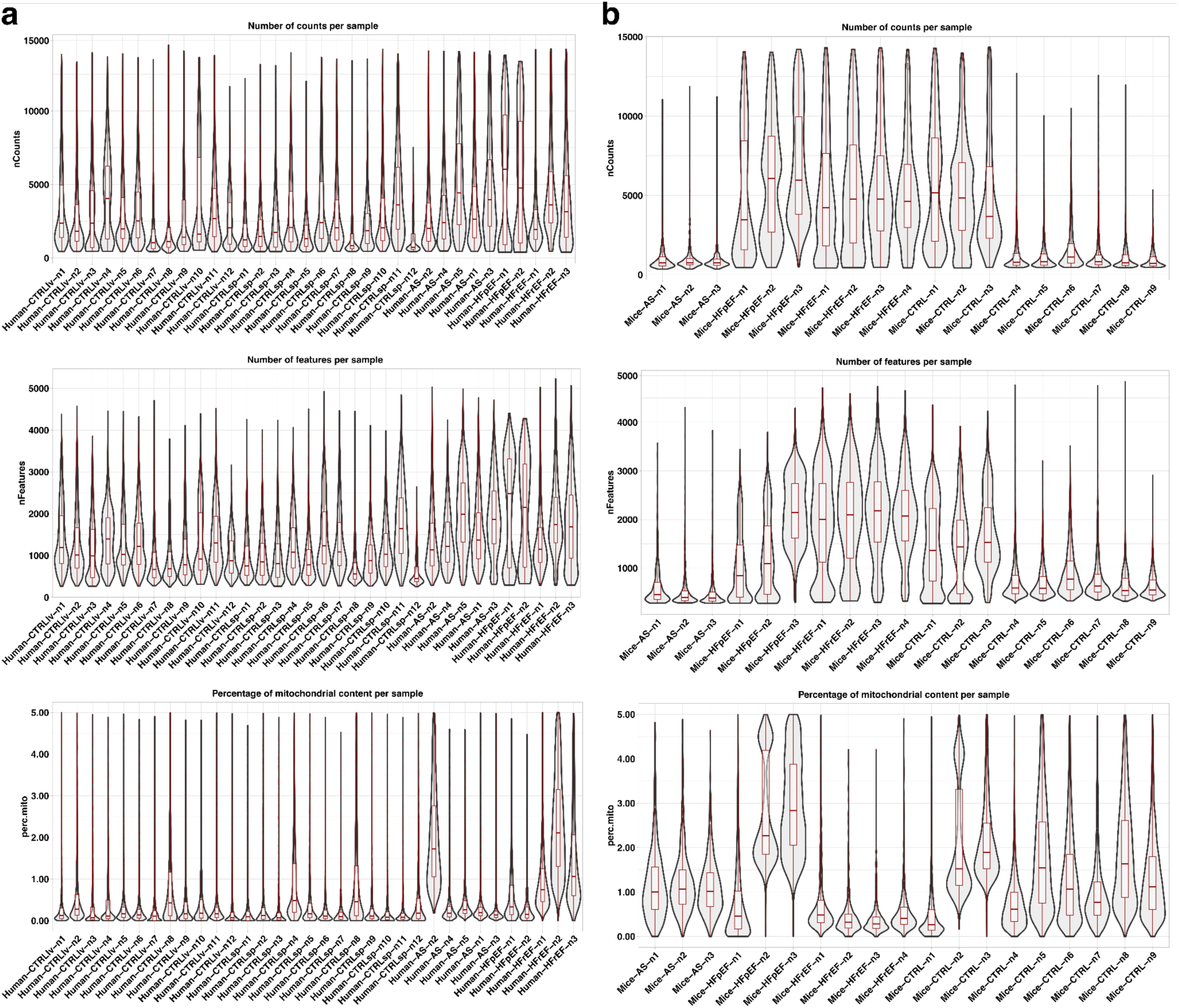
Quality measurements of snRNA-seq cells used for training, validation and test sets. **a-b**, Number of counts per sample, number of features per sample and percentage of mitochondrial content per sample for each heart failure state obtained in **a)** patients and **b)** mice models.

**Extended Data Fig.2:**
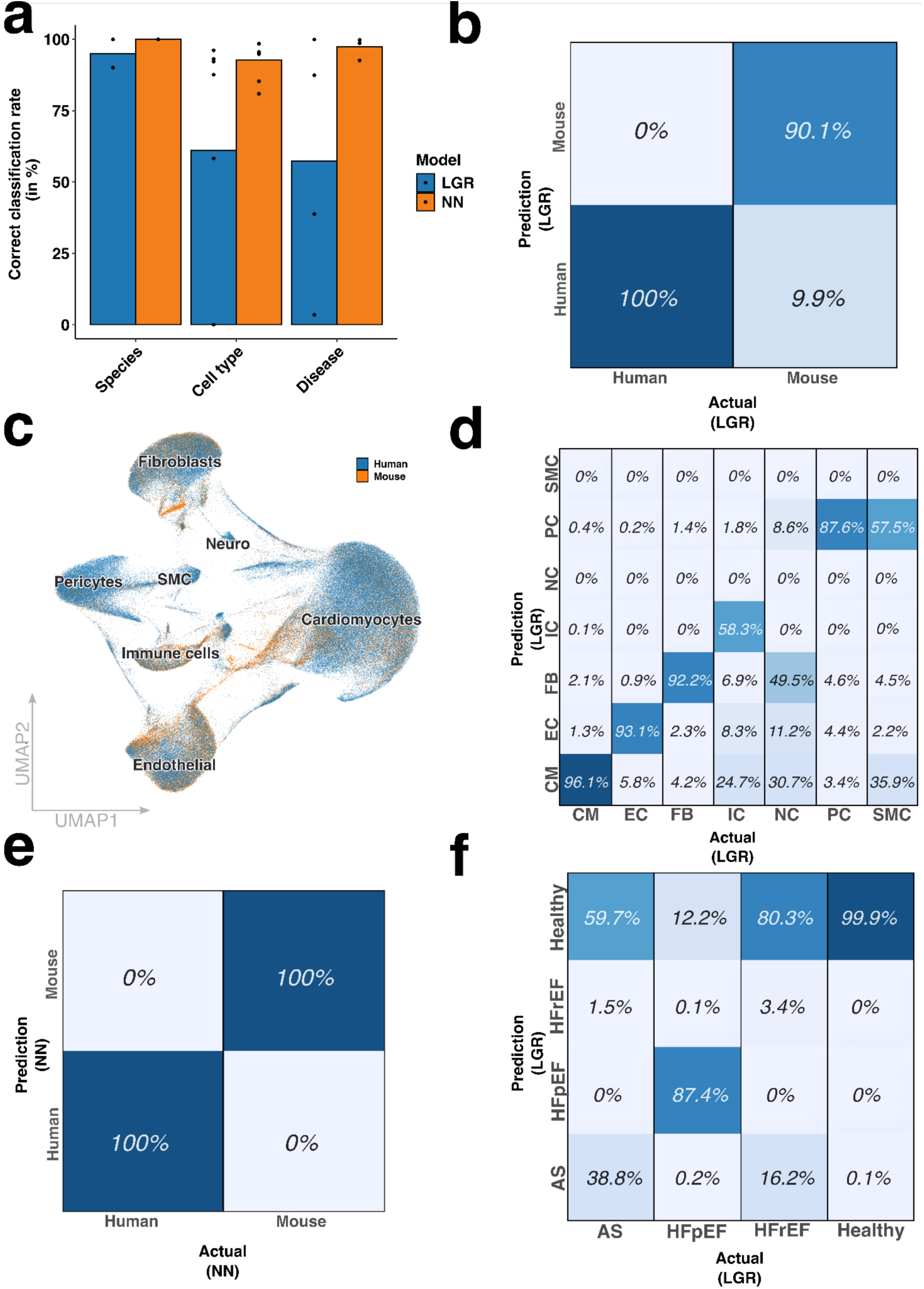
A logistic regression model can not capture the complexity of the data. **a,** Correct classification rates averaged for species, cell type and disease state. Calculations were made for the logistic regression model (LGR, blue) and for the neural network (NN, orange). **b,e** Confusion matrix calculated for the species in **b)** the logistic regression model and **e)** using the results of the neural network. **c**, Uniform manifold approximation and projection of cells showing the integration of human (blue) and mouse (orange) cells. **d,f** Confusion matrix calculations for evaluating the result of the logistic regression model regarding correctly classifying **d)** cell types and **f)** disease state.

**Extended Data Fig.3:**
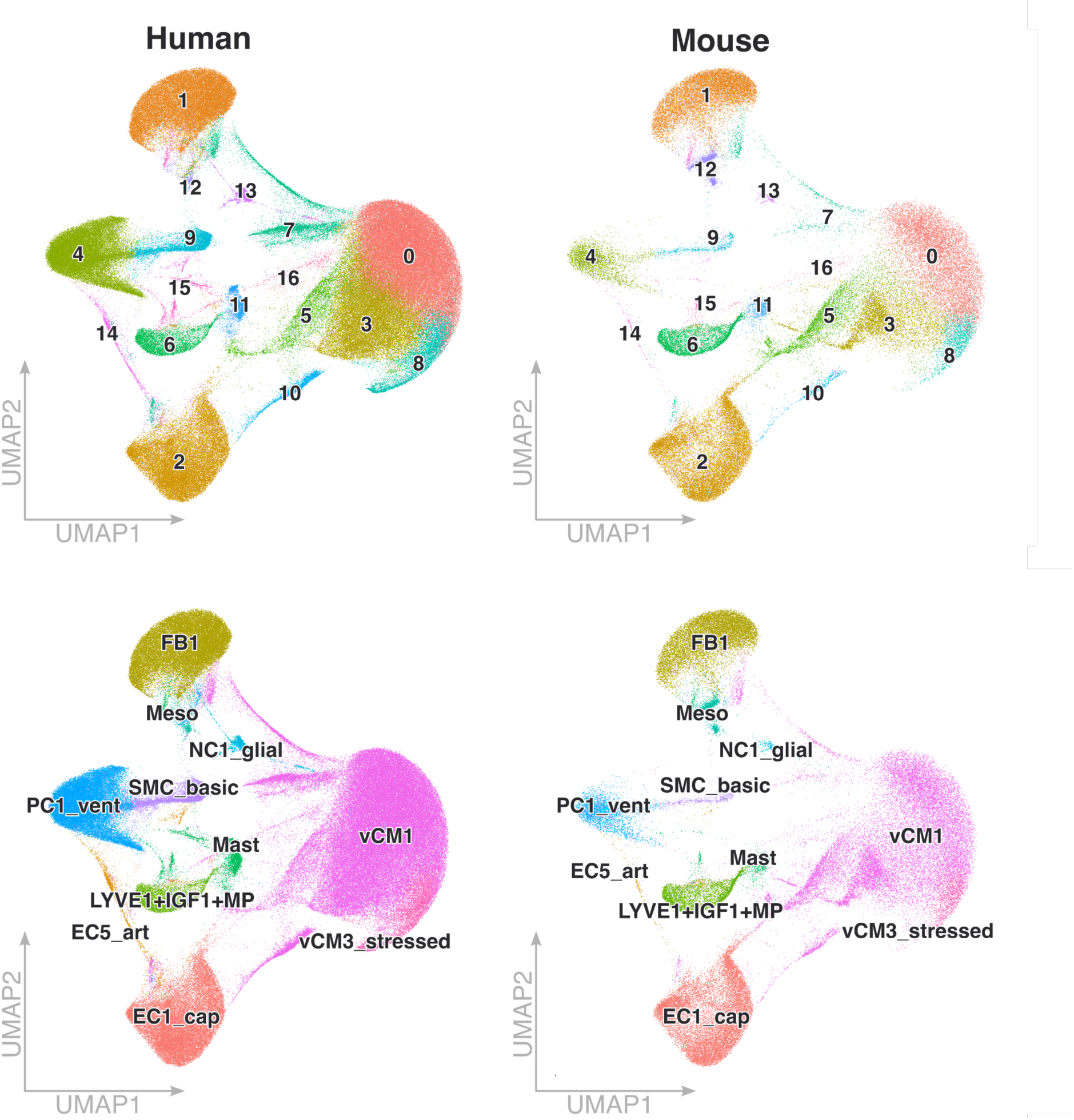
Automatic annotation of unsupervised clustering representing the data. Automatic annotation of the 16 identified clusters using CellTypist. The identified clusters belong to cardiomyocytes from the ventricle (vCM1, vCM3), capillary and arterial endothelial cells (EC1, EC5, respectively), fibroblasts (FB1), ventricular pericytes (PC1), smooth muscle cells (SMC), cells which show LYVE1 and IGF expression or are macrophages (LYVE1+IFG1+MP), neuronal cells (NC1), mesodermal cells (Meso) and mast cells (Mast).

**Extended Data Fig.4:**
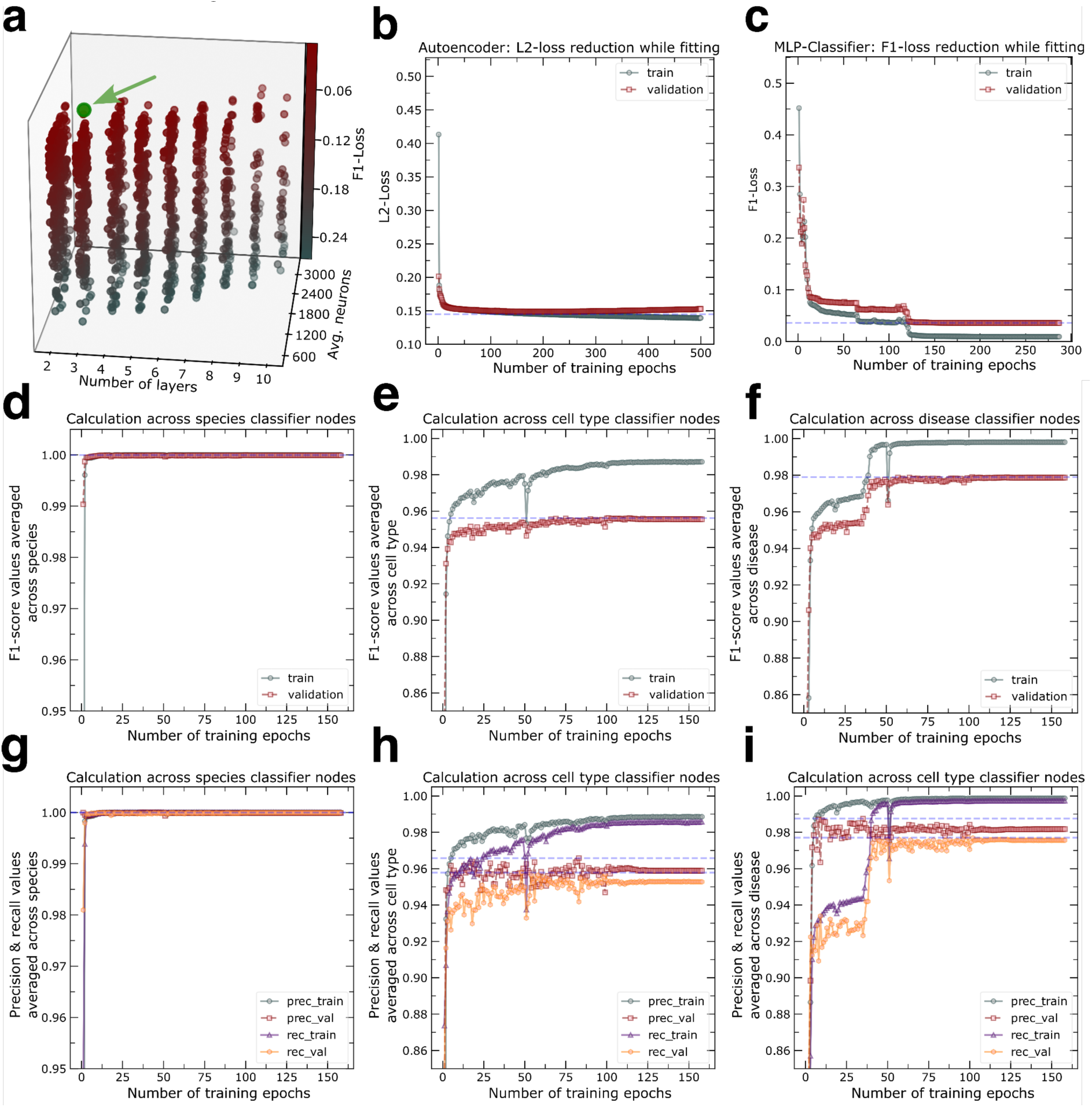
Learning curves of neural networks show little evidence of overfitting but high F1-scores. **a**, Three-dimensional scatter plot illustrating the F1-Loss in the MLP classifier layer, along with the corresponding number of neurons for each layer configurations tested during hyperparameter tuning. **b**, L2-loss calculations per epoch of the DAE considering the training data (blue) and the validation data (red). **c**, F1-loss calculations per epoch of the MLP classifier for the training data (blue) and for the validation data (red). **d-f,** Per epoch F1-score calculation during the classifiers fitting to the heart failure data averaged across **d)** species, **e)** cell type and **f)** disease state. **g-i,** Per epoch precision and recall calculations for the classifier while fitting averaged across **g)** species, **h)** cell type and **i)** disease state.

**Extended Data Fig. 5:**
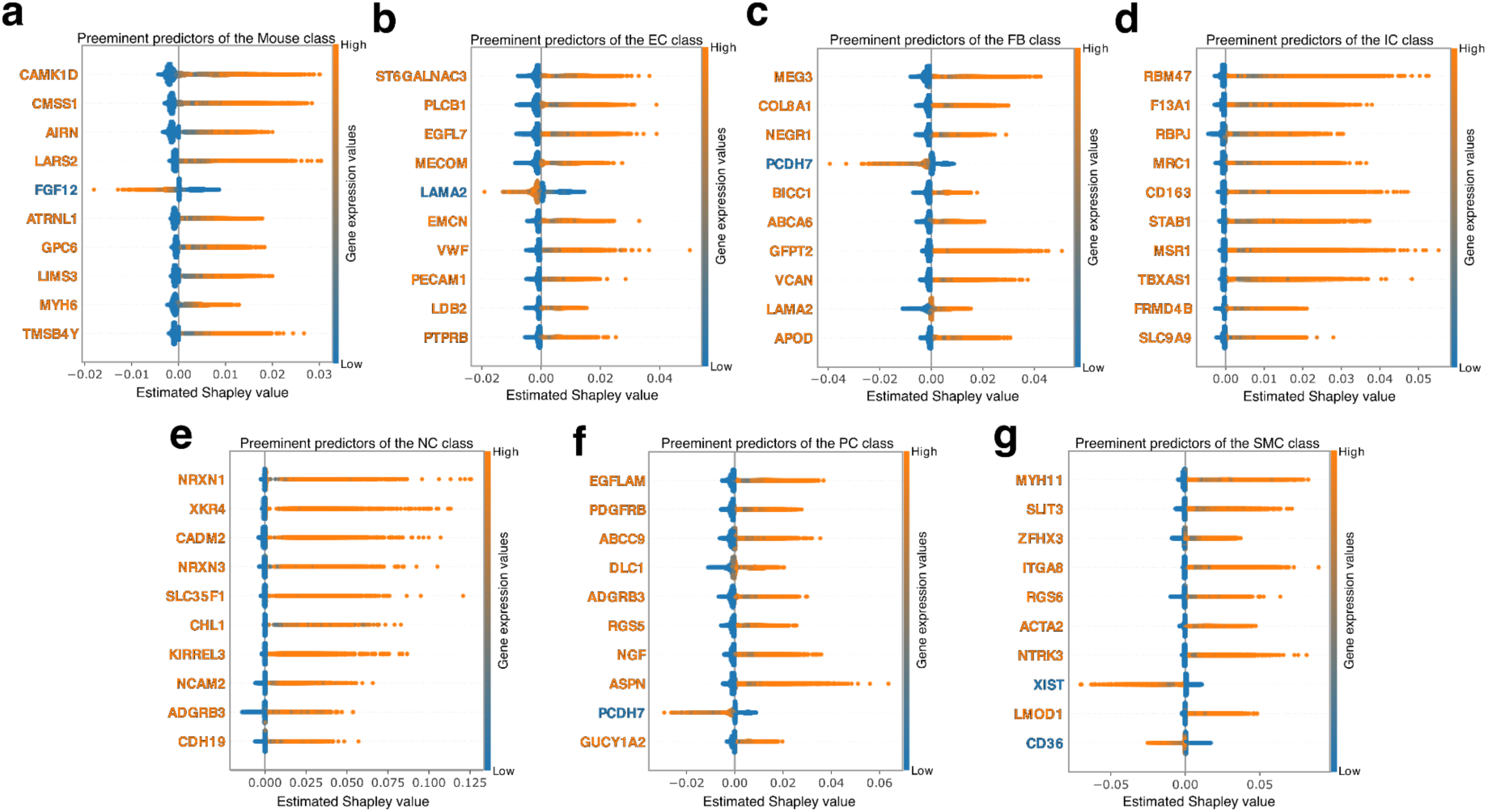
XAI analysis of the classifier nodes for mouse and the remaining cell types. **a-g**, Most influential predictors contributing to the **a)** mouse class, **b)** endothelial cell (EC) class, **c)** fibroblast (FB) class, **d)** immune cell (IC) class, **e)** neuronal cell (NC) class, **f)** pericyte (PC) class and **g)** smooth muscle cell (SMC) class. High expression values in the data are highlighted in orange while low expressions are blue. Depending on the Shapley values, the levels of expression can therefore be associated with a higher or lower prediction result for a given gene. An orange gene name indicates a positive correlation between expression and prediction for that class, while a blue gene name indicates a negative correlation.

**Extended Data Figure 6.**
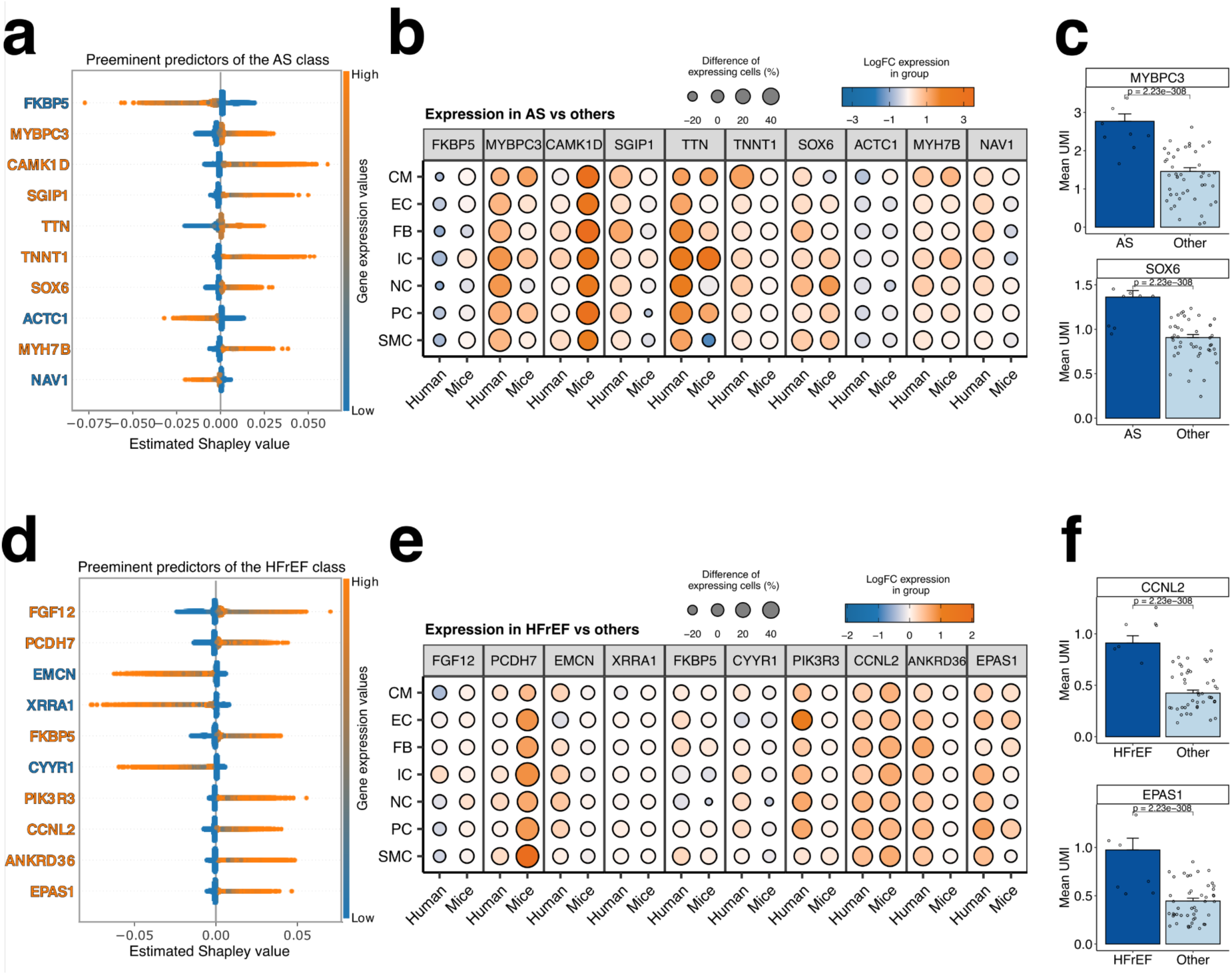
XAI analysis for heart failures caused by aortic stenosis (AS) and heart failure with reduced ejection fraction (HFrEF). **a,d,** Ten most contributing predictors visualized by a beeswarm plot for **a)** aortic stenosis and **d)** HFrEF cells. An orange gene name indicates a positive correlation between expression and prediction for that class, while a blue gene name indicates a negative correlation similar as in Figure 3. **b,e,** Dot plot illustrating the log fold change in gene expression and the percentage change of cells expressing the indicated genes. Panels represent the following **b)** AS predictors by cell type across human and mouse and **e)** HFrEF predictors by cell type and species. **c,f,** Averaged unique molecular identifiers (UMI) for two selected genes for **c)** aortic stenosis against the remaining conditions and **f)** HFrEF cells against the remaining conditions. Statistical significance was determined using the Wilcoxon test on single cell expressions.

